# Subunit-specific behavioral modulation of sensory tuning in the visual cortex

**DOI:** 10.1101/2025.09.04.674251

**Authors:** Julia M. Mayer, Wiktor F. Młynarski

## Abstract

Activity of sensory neurons is influenced not only by external stimuli but also by the animal’s behavioral state. It is well documented that behavior influences the general properties of neural activity, such as response gain. However, it is not known whether it could affect the sensory tuning of individual neurons in a more refined way and what the functional benefit of such nuanced modulation might be. Here, we investigate this in the mouse visual cortex using the data made available by the Allen Brain Observatory. Our analysis indicates that locomotion can modulate not only the gain of the entire neuronal response, but also more selectively control responses to specific stimuli. This modulation results in changes of neuronal tuning in different behavioral states. Using numerical simulations, we demonstrate that such patterns of gain modulation can multiplex behavioral information in sensory populations without compromising the accuracy of sensory coding. In that way, the visual cortex could instantiate an accurate, joint representation of sensory and movement-related signals and support computations that simultaneously require both types of information.

## INTRODUCTION

With the advent of neural recordings in behaving animals, it became clear that sensory neurons are continuously modulated by multiple non-sensory signals [1]. For example, neurons in the primary visual cortex (V1) of rodents can be modulated by spontaneous behavior and body movements [2, 3], location in the environment [4], head and eye position [5], and internal states such as arousal [6] and attention [7]. Modulation of sensory neurons by animal movement raises general questions about sensory coding during active behavior.

A behavioral state that exerts a particularly strong impact on sensory activity is locomotion speed [8]. Neurons in the visual system can dramatically modulate their gain depending on the movement of the animal [9–12]. In it’s simplest form, this modulation is applied to all stimuli identically (Fig.1a, left panel). This means that at any given running speed, the sensory tuning of the neuron is characterized by the same tuning curve, which is scaled by a speed-dependent factor. In direction- and orientation-selective cells, this mechanism modulates the amplitude of the uni-modal or bimodal tuning curves respectively (Fig.1a, right panel).

**Figure 1:**
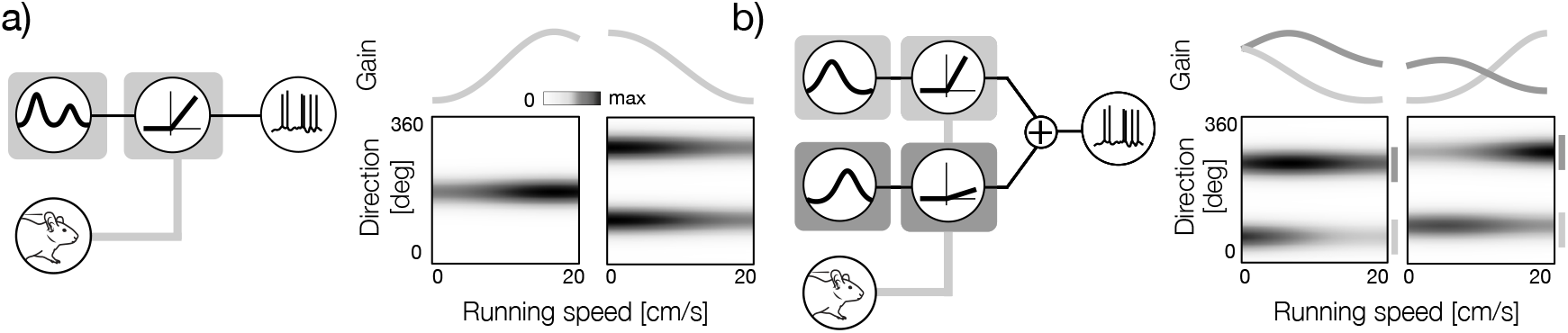
Multiplexing sensory and behavioral signals with gain modulation. **a)** Left: a functional neuron model where the gain of a sensory tuning curve is modulated by the running speed. The modulation is applied identically to the entire tuning curve. Right: two example direction-speed tuning curves corresponding to the model in the left. A direction-selective (left) and orientation-selective (right). The behavioral modulator as a function of the running speed is plotted above tuning curves. **b)** Left: a functional neuron model where the running speed independently modulates the gain of the different parts of the tuning curve (subunits). Right: two example direction-speed tuning curves corresponding to the model in the left. Sensory subunits are marked with vertical lines in different shades of gray to the right of the tuning curve. Their modulators are plotted above the tuning curves in the corresponding color.

The input-output relationship implemented by sensory neurons is, however, often much more complex. Even individual cells can perform complex computations that involve multiple stimulus features and sequences of transformations [13]. These individual stimulus features are frequently referred to as “subunits [14–20]. Individual subunits can be associated with dedicated nonlinear transformations that control their gain [15]. This complexity of tuning raises the possibility of a more nuanced form of behavioral modulation in the visual system. Hypothetically, if the stimulus transformation implemented by a neuron operates on multiple distinct subunits, each of the subunits could be modulated independently by behavior (Fig.1b, left panel). Neurons characterized by this functional architecture could vary their sensory tuning at different locomotion speeds (Fig.1b, right panel). This modulation could potentially allow them to implement complex computations with sensory and behavioral signals. The existence of such neurons has not yet been confirmed.

Here, we analyze neural activity in the mouse visual cortex and provide evidence that neurons indeed modify their tuning depending on the animal’s speed. We computed two-dimensional tuning curves that characterize how visual neurons are modulated by running speed. We found that the tuning of sensory neurons in the mouse visual cortex is modulated by the behavioral state. This modulation takes a specific form: running speed can differentially adjust the gain of fragments of the tuning curve, which we refer to as functional subunits of a neuron. Using a sparse decomposition of orientation-velocity tuning curves, we extracted the shape of subunit gain as a function of the animal running speed. We found a diversity of gain modulation patterns used by cortical neurons to dynamically change the weights of sensory inputs at different locomotion states.

Using numerical simulations, we demonstrate that behavioral modulation can have a positive effect on stimulus encoding, even if the features combined by the neuron are modulated differentially by behavior. Neurons that implement such dynamic tuning retain information about the modulating variable, thus instantiating an accurate joint representation of stimuli and the behavioral state. We speculate that such representations could support computations that require sensory and behavioral signals, and that they could be easily implemented by combining the inputs of neurons with simpler forms of tuning. Overall, our results suggest that the mouse visual cortex instantiates an accurate multiplexed representation of the behavioral state and the sensory environment.

## RESULTS

### Estimation of joint tuning to the stimulus direction and running speed in the mouse visual cortex

Our aim is to test whether the sensory tuning of neurons can be modulated by the behavioral state. To this end, we explored two-photon calcium imaging recordings of the mouse visual cortex made available by the Allen Brain Institute [21].

The first step of our analysis was to determine the simultaneous tuning of neurons in the mouse visual cortex to a stimulus direction and running speed, important sensory and behavioral variables. The Allen Brain Institute recordings were performed in head-fixed behaving mice whose running speed was acquired simultaneously with neural activity (Fig. 2a). Mice were exposed to drifting grating stimuli at different temporal frequencies and movement directions (see Methods). To understand how sensory tuning interacts with behavior, we needed to estimate the tuning at different running speeds. This presented a challenge because individual animals had different behavioral preferences and generated very different distributions of running speed. Since our aim was to pool and compare the data recorded in multiple sessions and animals, we could not rely on quantile binning as suggested before [9, 10]. Instead, we used 8 bins linearly spaced between 0 and 20 cm/s.

**Figure 2:**
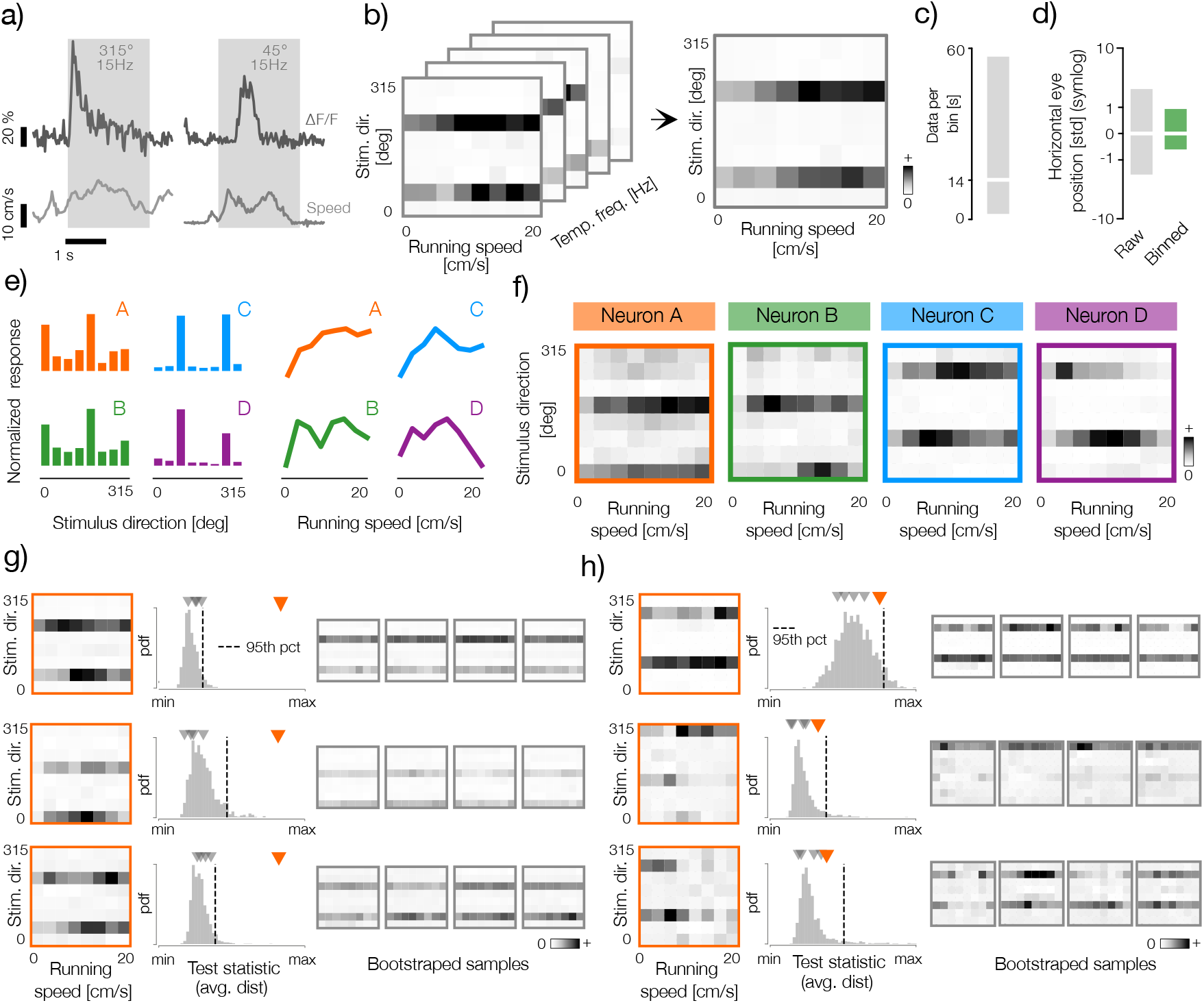
Analysis of tuning to stimulus direction and running speed in the mouse visual cortex. **a)** Example traces of the calcium imaging data (dF/F) and of mouse running speed obtained from the Allen Brain Observatory (top, dark gray lines and bottom, light gray lines respectively). Gray rectangles mark duration of the stimulus whose orientation and temporal frequency are marked with gray text. **b)** Left panel: example three-dimensional (running speed - stimulus direction - stimulus temporal frequency) tuning curve of a neuron computed using thresholded dF/F signal (see Methods). Right panel: a two-dimensional (running speed - stimulus direction) tuning curve obtained by averaging the three-dimensional tuning curve on the left across the temporal frequency dimension. **c)** Distribution of seconds of data per bin in all analyzed datasets. The horizontal white line denotes the average (14 s) and the gray rectangle denotes the interval spanned by 5th and 95th percentiles of the distribution. **d)** Distributions of the horizontal eye positions pooled across raw recordings (left, gray bar) and averaged within bins of the orientation-speed tuning curves (right, green bar). White lines denote the mean (0 deg) and rectangles denote the interval spanned by 5th and 95th percentiles. Data for each animal was standardized independently via z-scoring. The scale of the plot is symlog (i.e. sign *x* log_10_( |*x*|). The eye position data was not available for all analyzed datasets. **e)** Left panel: orientation tuning curves of four example neurons. Right panel: running-speed tuning curves of the same neurons as in the left panel (marked with corresponding letter and color). **f)** Joint orientation-speed tuning curves of four example neurons marked with corresponding letters and colors in e. **g)** Example neurons that pass the bootstrap test of velocity modulation. Left column: orientation-speed tuning curves of three example neurons (orange frames). Middle column: distributions of the test statistic of the modulation test (see Methods). Vertical dashed line denotes the significance threshold equal to the 95th percentile of the statistic distribution. Orange and gray triangles denote respectively the values of the test statistic of the original tuning curve and five example bootstrapped samples visualized in the right column. Right column: five example bootstrapped orientation-speed tuning curves. **h)** Example neurons that do not pass the bootstrap test of velocity modulation. Panel is analogous to g).

Using this binning scheme, we computed three-dimensional tensors (“tuning curves”) for each neuron by pooling and averaging the normalized dF/F signal for each bin specified by the stimulus orientation, its temporal frequency, and the running speed of the animal (Fig. 2b, left panel). Each three-dimensional tensor was then averaged across the temporal frequency axis, generating a two-dimensional direction-speed matrix, which we refer to as a direction-speed tuning curve (Fig.2b, right panel). To ensure that the tuning curves were computed using a comparable set of stimuli, we selected only the data sets with minimal differences in stimulus frequencies presented in each direction and running speed (see Methods). In this way, we aggregated a reasonable amount of unbiased data for each direction-speed bin (average 14 seconds, Fig.2c). This pooling scheme ensured that we averaged across data recorded at more eye positions within each bin, which effectively reduced eye position variability across all bins of the tuning curve (Fig. 2d, gray and green bars, respectively).

Since the focus of our study is on adjustments of sensory code by behavioral variables, we limited our analysis only to neurons that are significantly tuned to stimulus direction, as measured by the sharpness of their tuning to stimulus direction (see Methods). Tuning of four example neurons to the stimulus direction and the running speed is shown in Fig.2e (left and right panels, respectively). Pairs of neurons denoted by A, B as well as C, D have similar sensory preference and respond with comparable strength to stimuli with the same direction. Behavioral tuning does not appear to be related to the sensory preferences of these cells. Here, different pairs of cells (A, C and B, D) respond similarly to the running speed of the animal. If these example neurons encoded behavioral and sensory signals independently of each other, their tuning could be expressed in full using these two one-dimensional tuning curves. However, inspection of the joint tuning to running speed and stimulus direction reveals that this is not the case (Fig.2f). In each example neuron, sensory tuning visibly varies across the considered range of running speed. Although neurons do not entirely shift their orientation-tuning curves, they seemingly modulate the gain of their responses to preferred stimuli in different behavioral states. The gain modulation patterns are distinct in each cell, suggesting that they could represent different joint features of sensory inputs and behavior.

The example neurons displayed in Fig.2e-f clearly reveal different patterns of joint direction-speed tuning. However, their tuning curves were computed using different amounts of data, which itself are variable and noisy. To ensure that the computed tuning curves reflect preferences of individual neurons, and not biases in the data distribution or noise, we performed a bootstrap analysis (Fig.2g-h). For each neuron, we randomly shuffled the data across running speed values and computed estimates of the direction-speed tuning curves (Fig.2g-h, right columns). This bootstrapping scheme preserved the sensory tuning of each cell (both to the stimulus direction and temporal frequency) and the amount of data per bin, modifying only the impact of the running speed on the response gain (see Methods for details). The majority of neurons were significantly modulated by behavior, exceeding the chance level established via bootstrapping(Fig.2g). Neurons that did not pass the significance test (e.g. examples in Fig.2h) were excluded from further analysis.

### Sparse decomposition of tuning curves reveals subtle effects of running speed on sensory tuning

The shape of the orientation-speed tuning curves in the mouse visual cortex suggests that neurons may change their sensory tuning at different running speeds. This dynamic tuning can be functionally characterized by two direction-selective subunits (i.e. parts of the tuning curve tuned to stimuli separated by 180 degrees), whose gain is differentially modulated by running speed (Fig.3a). We note that is only a quantitative characterization of neuronal tuning, which could be implemented by different neuronal mechanisms (see Fig.9).

To separate the impact of movement on these individual subunits, we fitted each tuning curve with the sparse-decomposition model (Fig.3b). The components of the model correspond directly to elements of the functional architecture in Fig.3b. The model represents the average neural activity summarized in a tuning curve as a sum of basis functions that correspond to sensory subunits tuned to a specific directions of visual motion (Fig.3b, second and forth panels from the left). The row of each subunit basis function is point-wise multiplied by a corresponding velocity modulator, i.e. the curve that describes how the gain of that subunit changes with the speed of the animal (Fig.3b, third and fifth panels from the left). During fitting, we placed a separate sparse penalty on individual subunits and on modulators, which increased the robustness and quality of the fits (see Methods). We excluded from further analysis cells that were not significantly well fitted by the sparse decomposition model, which we assessed by bootstrap testing (see Methods). The tuning curves of the retained neurons were well approximated by the model fits, with the average fit quality equal to 8 dB SNR (Fig. 3c).

**Figure 3:**
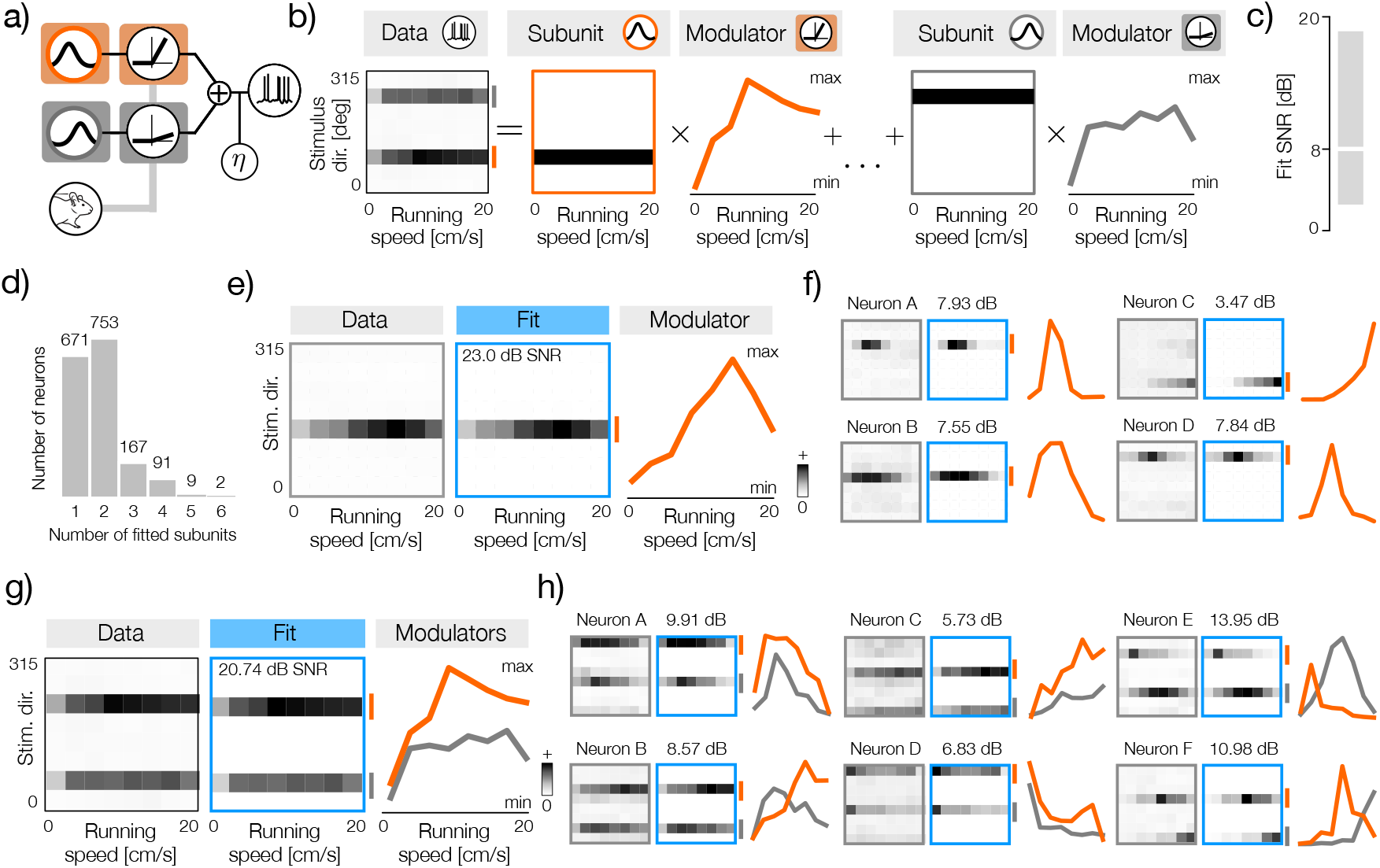
Sparse decomposition of direction-speed tuning curves. **a)** Conceptual schematic of a neuron where two sensory subunits (orange and gray circles) are modulated independently by the running speed of the animal. **b)** Schematic of the statistical model that decomposes stimulus-speed tuning curves (left column) into a sum of subunit basis functions (second and fourth panels from the left) multiplied by speed-controlled gain modulators (third panel from the left and right panel). The subunit basis functions determine sensory tuning of a neuron, while modulators determine modulation of each subunit by the running speed of the animal. **c)** Distribution of the SNR values of model fits to stimulus-speed tuning curves. The horizontal white line denotes the average (8 dB SNR) and the gray rectangle denotes the interval spanned by 5th and 95th percentiles of the distribution. **d)** Histogram of the number of subunits fitted to each neuron. Note that not a single neuron had 7 or 8 significant subunits. **e)** Left panel: example stimulus-speed tuning curve of a direction-selective neuron with a single sensory subunit. Middle panel: reconstruction of the tuning panel from the model fit. The single fitted sensory subunit is marked with a short vertical orange line on the right side. The SNR of the fit is equal to 23 dB. Right panel: speed modulator of the direction-selective subunit. **f)** Four example direction-selective neurons (gray frames) together with their respective fits (blue frames) marked with fitted subunits (short, vertical orange lines on the right side of each blue frame) and speed-modulators (orange lines). **g)** Same as e but for an example direction-selective neuron with two sensory subunits. The subunits and their modulators are now marked with gray and orange lines. **h)** Same as f but for six example orientation-selective neurons.

The parameter of the fitted model identified the selectivity for direction and orientation in the mouse visual cortex, consistent with previous findings [22, 23]. The largest number of neurons had 2 significant subunits (N=753, see Methods for the definition of significant subunits). If two subunits differ by 180 degrees in their sensory preference, they instantiate orientation tuning, which was predominantly the case (e.g. Fig.3g). The second largest group of neurons (N=671) was best fitted by a single significant subunit (e.g. Fig.3e), which corresponds to direction selectivity. In other neurons, additional significant subunits typically modeled broadened sensory tuning. We therefore concentrate our analysis on direction and orientation selective cells with either 1 or 2 significant subunits, respectively.

The sparse decomposition model successfully extracted speed modulators from the tuning curves of direction-selective neurons (Fig.3f). Estimated modulators take different forms, e.g., increasing (Fig.3f, neuron C) or non-monotonic and preferring a specific speed (Fig.3f, neurons A, B, D). These findings are in agreement with previous studies that reported a diversity of speed tuning in the visual system of a mouse [9–11]

In orientation-tuned neurons, we obtain two modulators, one for each direction-selective subunit. (Fig.3h). Interestingly, the differences between modulators of individual subunits can take various forms. Individual subunit modulators can increase or decrease at different rates, instantiating differential sensory tuning at various running speeds (Fig.3, neurons C, D). Individual modulators are also often non-monotonic and can either peak at similar running speeds (Fig.3h, neuron A) or, in a dramatic demonstration of the shift in sensory tuning, amplify responses at different speeds of locomotion (Fig.3h, neurons B, E, F).

### Bootstrap analysis reveals influence of running speed on tuning of sensory neurons

Following the extraction of speed-modulators that characterize gain changes of individual subunits, we can now quantify whether these can be significantly different in single neurons and how often such differences occur in the population. We perform this quantification using a bootstrapping procedure that occurs in three steps. First, for each neuron, we use a shuffling scheme to simulate a population of neurons with identical sensory tuning preferences but with shared subunit modulation induced by running speed. Second, for each sample bootstrapped in this way, we compute a test statistic that quantifies similarity of two modulators. Finally, we compare the value of the test statistic derived from the real tuning curve to the distribution of the bootstrapped samples. We describe these steps in more detail below.

As before, for each neuron, we randomly shuffled the data across running speed values, removing the effect of speed modulation. We then multiplied neural responses to preferred stimulus directions by a common modulator, which was computed by averaging two significant modulator fits recovered by the sparse decomposition model. This bootstrapping scheme preserved the sensory tuning of each cell to the stimulus direction and temporal frequency, as well as the amount of data per bin. The influence of the running speed was, however, replaced by a shared multiplicative modulation of significant subunits (Fig.4a-b). For each neuron, we generated 1000 such randomized samples.

If both subunits of an orientation selective neuron are modulated jointly by running speed, their modulators should be accurately represented by a single vector. This intuition guided our design of an appropriate test-statistic. For each real tuning curve and a bootstrapped sample, we optimized a correlation-maximizing fit, i.e., a curve that maximizes Pearson’s correlation with both modulators on average (Fig.4c-d, black dashed lines). We denote the value of this correlation for each real tuning curve as 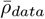, for each bootstrapped sample as 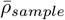, and we take it to be the basis for our comparison. Neurons whose modulators were represented by a single fit as good as their corresponding bootstrap samples (e.g. Fig.4d) most likely do not change their tuning with running speed. On the other hand, neurons for which a single fit was correlated with individual modulators much more weakly than in the simulated bootstrap samples were likely modulating their sensory tuning across locomotion states (e.g. Fig.4c). We derived the significance threshold from the test statistic 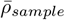 computed for each of the 1000 tuning curves with simulated shared modulation. If the statistic 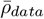 lies below the threshold, the neuron can be considered to significantly change it’s sensory tuning with the running speed (Fig. 4 e). If the data value exceeds the threshold, the neuron’s subunits can be explained by a common modulatory input that does not modify the sensory tuning (Fig.4 f). A summary of the test statistic distribution for a random subset of 150 neurons is visualized in Fig.4g.

**Figure 4:**
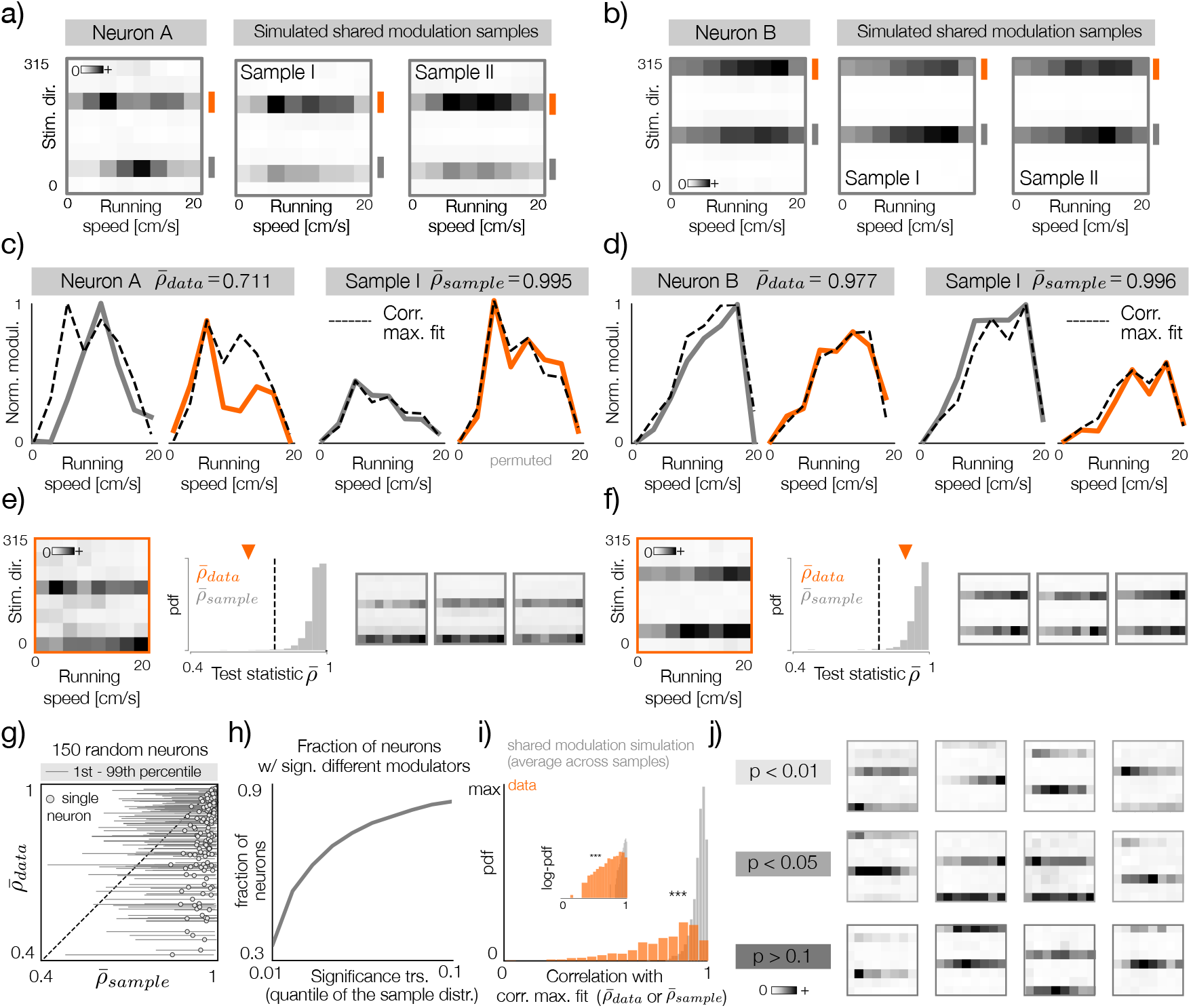
Quantification of differential modulation of sensory subunits. **a)** Left: tuning curve of an example neuron with differentially modulated subunits. Right: two bootstrap samples with simulated shared modulation corresponding to the neuron on the left. **b)** Same as a but for an example neuron with shared modulation between subunits. **c)** Left: modulators fitted to the tuning curve in a (orange and gray lines correspond to marks in panel a). Dashed black line denotes the correlation maximizing fit to both modulators. Note that the fitted curve is identical for both modulators up to a scaling and a shift. The average correlation of the fit with the two modulators 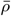 is equal to 0.711. Right: the same as on the left but for bootstrap sample I depicted in panel a. **b)** Same as c, but for the example neuron displayed in b. **e)** Left: example tuning curve of a neuron with differently modulated subunits. Middle: a null distribution of the test-statistic 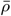 i.e. the average correlation of the correlation-maximizing fit with both modulators, derived from 1000 bootstrap samples with simulated shared modulation. The vertical dashed line denotes significance threshold at *p* = 0.01 and the orange triangle the value of the test statistic for the tuning curve on the left. Right: three example bootstrap samples with simulated shared modulation. **f)** Same as e, but of an example neuron whose subunits are not significantly differently modulated. **g)** Visualization of the test statistic distribution for 150 randomly selected neurons. Gray circles denote the value of the test statistic 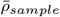 averaged across all bootstrap samples (x-axis) and the value for the data 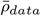 (y-axis). Thin-gray lines denote the range between 1st and 99th percentile. **h)** Fraction of neurons that pass the significance threshold of the test for differential modulation of sensory subunits as a function of the threshold value. The threshold is computed as a quantile of the null distribution obtained for each neuron individually via bootstrapping. 35 percent of neurons pass the significance threshold at *p* = 0.01. **i)** Histogram of the test statistic for the data (orange) and the average across bootstrap samples for each neuron (gray). The distributions are significantly different (KS-test, p-value *<* 0.001). Inset depicts the same distribution on the log-probability scale. **j)** Example tuning curves that pass the significance threshold at p-value thresholds of 0.01, 0.05 (top and middle rows respectively) and that do not pass the threshold of 0.1 (bottom row).

At the significance threshold p=0.01, approximately 35% of the population reveals significant modulation of sensory tuning with running speed. This proportion increases rapidly with the significance threshold, up to 90% at p=0.1 (Fig.4h). The pronounced difference between the real and simulated tuning curves is clearly visible in the distributions of the test statistic 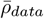 and 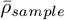 averaged across all bootstrap samples (Fig.4i).

The significance of modulator differences depends on three factors: the existence and strength of the underlying biological process, the distribution of data across direction-speed bins, and noise (Fig.4j). Tuning curves that achieve significance at progressively increasing thresholds range from clearly distinct subunit modulation patterns (p < 0.01) to less distinct and noisier patterns (0.01 < p < 0.05). Tuning curves that do not pass the significance test even at a relatively high p-value (p > 0.1) have nearly identical modulators, differing only in the sampling noise or relative subunit strengths. In our interpretation, these results indicate a continuum of effects that running speed exerts on sensory tuning. At the very least, a considerable fraction of the population (approximately one third) modulates it’s sensory tuning across behavioral states. In consecutive analyzes, we therefore consider all orientation-selective neurons jointly and do not separate them according to the test outcome. Understanding the extent and strength of this effect in detail requires additional data and new experiments.

### Analysis of speed induced gain modulation in orientation-selective neurons

In a large majority of two-subunit neurons (83%), individual subunits encode stimulus directions separated by 180 degrees (Fig.5a, right panel). The joint distribution of stimulus directions preferred by individual subunits does not reveal any systematic bias (Fig.5a, left panel).

Orientation-selective cells combine individual subunits asymmetrically, with unequal weights. We quantified this effect by averaging the modulators of each subunit and taking the ratio of the stronger to the weaker modulators. Although the most likely ratio is equal to 1, the average ratio of the modulator strengths of the subunits is 3.36 and spans a wide range, as indicated by the distribution of the ratio logarithms (Fig.5b). We note that the asymmetry of subunit strengths is a signature of the neuron differentiating between two directions of visual motion separated by 180 degrees, which could, in principle, be inherited from behavioral modulation.

The similarity of subunit modulators in the same neuron can be easily quantified using correlation. The correlations of the modulator pairs are broadly distributed, covering a wide range from highly positive to highly negative values (Fig.5c, orange bars). Although the distribution is broad, it is clearly skewed towards more positive correlations, indicating that many modulator pairs are similar. To ensure that this pattern is not random, we computed correlations between modulators of bootstrap tuning curves where the effect of speed modulation has been removed (Fig.5c, gray bars; see Fig. 2g-h). As an additional control, we computed correlations of modulators in cells with simulated shared gain modulation (Fig.5c, blue bars; see Fig. 4a-b). The data distribution is significantly different from these two controls (KS-test, p-value ≤ 0.001), suggesting that the observed distribution can not be explained entirely either by random noise nor by a shared modulatory signal.

To understand how individual modulators are related to the behavioral state, we computed their correlations with the running speed of the animal (Fig.5d, orange bars). In stronger and weaker subunits (Fig.5d, orange bars, left and right panel, respectively), the correlations with speed are statistically indistinguishable (KS test, p-value = 0.24). In both cases, the values span a wide range of correlations, with the mode being close to a perfect correlation (i.e. *ρ* = 1), which corresponds to a monotonic increase in gain with running speed. The correlation distributions for individual modulators are also significantly different from the bootstrap samples without modulation (Fig.5d, gray bars), which further strengthens the notion that both modulators reflect significant biological processes and not noise. At the same time, the data is, as expected, statistically indistinguishable from the distribution of correlations with speed in bootstrap samples with simulated shared modulation (Fig. 5d, blue bars, p-value > 0.1).

**Figure 5:**
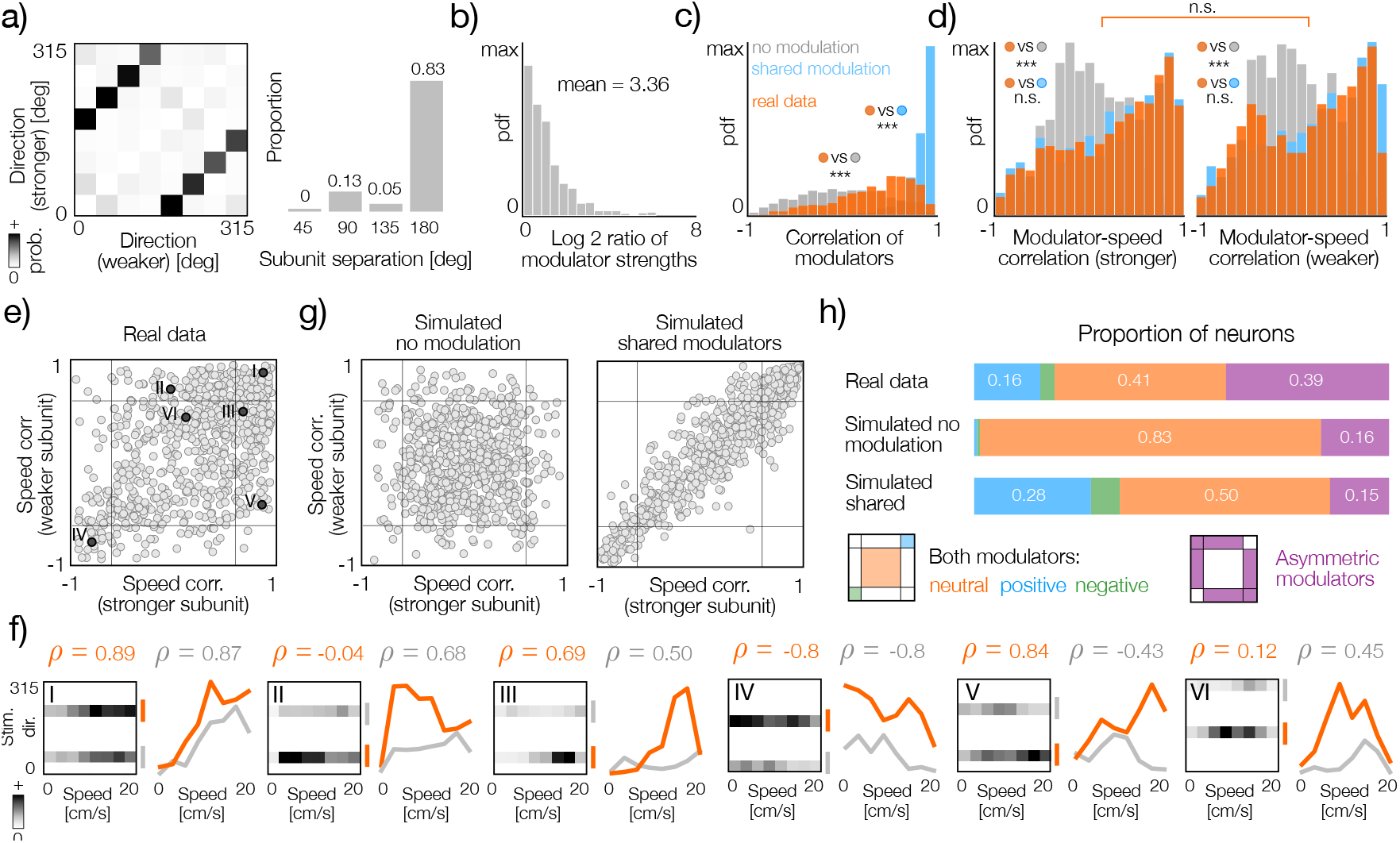
Behavioral modulation of orientation-selective cells. **a)** Left panel: joint distribution of stimulus directions preferred by subunits in two-subunit neurons. Right panel: histogram of angular separation (distance) between stimuli preferred by individual subunits in two-subunit neurons. **b)** Histogram of log-ratios of modulator strength (stronger to weaker). The mean equal to 3.36 was computed before logarithmic transform. **c)** Histograms of correlations between modulators in two-subunit neurons. Orange, grey and blue correspond to the original data, bootstrap tuning curves without modulation and bootstrap tuning curves with shared modulation respectively. Data distribution is significantly different from both simulated datasets (KS-test, both p-values ¡ 0.001). **d)** Histograms of correlations between modulators and running speed in stronger and weaker subunits (left and right panels respectively). Orange, grey and blue correspond to the original data, bootstrap tuning curves without modulation and bootstrap tuning curves with shared modulation respectively. Correlation distributions are significantly different from bootstrap without modulation in both panels (KS-test, p-value *<* 0.001) and not significantly different from bootstrap with shared modulation (KS-test, p-value *>* 0.1). Correlation distributions for stronger and weaker modulators are not significantly different (KS-test, p-value=0.24). **e)** Scatter plot of modulator correlations with running speed. Each gray dot corresponds to a single neuron. Thin black lines within the plot denote 5th (left and bottom line) and 95th (right and upper line) percentiles of modulator-speed correlations computed with bootstrap samples without modulation. Darker points with Roman numerals correspond to identically labeled tuning curves in panel f. **f)** Example direction-speed tuning curves (grayscale heatmaps) and corresponding fitted modulators of stronger and weaker subunits (orange and gray lines respectively). Correlations of the stronger and weaker modulator with the running speed are denoted as *ρ* in orange and gray respectively. Roman numerals correspond to points marked in panel e. **g)** The same as e but for randomly permuted modulators (left) and simulated joint modulators (right). **h)** Proportions of cells in different regions of the correlation plane for real data (top row), bootstrap samples without modulation (second row from the top) and bootstrap samples with simulated joint modulators (third row from the top). Color legend of areas on the correlation plane (as in e-f) is depicted in the bottom row.

Distributions of modulator correlations with running speed characterize behavioral influences on individual subunits but not on the tuning of entire neurons. Changes in neuronal tuning induced by running speed are reflected in the joint relationship of the speed-modulator correlations visualized in Fig.5e. Each neuron is characterized by a point on the plane, whose horizontal and vertical axes correspond to the weaker and stronger subunits, respectively. To provide a reference and quantify joint-modulation patterns, we mark the 5th and 95th percentiles of speed correlations in bootstrap samples without modulation (Fig.5e left and bottom, right and top black lines, respectively). The scatter plot indicates that the speed correlation of one of the modulators does not necessarily determine that of the other. If one of the modulators is strongly correlated with speed, there is a broad range of possible correlations for the second modulator. We visualize example neurons corresponding to different speed correlation patterns in Fig.5f.

To ensure that the joint distribution of modulator speed correlations is not random and to interpret it more easily, we used two simulated joint distributions of modulator speed correlations (Fig.5g). First, we analyzed bootstrap samples without speed modulation (see Fig.2g-h). This resampling removes the significant speed-correlation of individual modulators, as well as their relationship with the other modulator in the same neuron (Fig.5g, left panel). Second, we analyzed bootstrap samples with simulated shared modulation (Fig.5g, right panel). In this way, we simulated a scenario in which the subunits of each neuron are jointly modulated by the same process, and any differences between modulator fits are due to random factors in the data. Both simulated distributions visibly differ from the data.

To quantify these differences, we computed the fraction of neurons in different areas of the correlation plane for the real data and the simulated datasets (Fig.5h). We divided the correlation plane into four areas. The first three areas correspond to modulator pairs that are simultaneously strongly positively correlated with speed, strongly negatively correlated with speed, or neutral; i.e., their correlation magnitude is not significant (Fig.5h, blue, green, and orange areas, respectively). We note that if both modulators are “neutral” with respect to running speed, they can be quite different from each other and modify the sensory tuning of a neuron at different running speeds. The fourth area corresponds to pairs of asymmetric modulators, where one modulator is strongly related to running speed and the other is not (Fig.5h, bottom row, purple area). The distribution of neurons in different regions of the correlation plane is clearly different between the real data and both simulated datasets (Fig.5h, top three rows). In particular, the number of asymmetric modulators, where each subunit is differently affected by running speed, is more than twice as high as in the bootstrapped datasets. These results further suggest that behavior can differentially modulate individual subunits, thereby adjusting sensory representation in different behavioral states. In the supplemental information, we describe additional control analyzes computed with alternative ways of simulating shared modulators (Suppl. Fig. 1).

### Clustering reveals diversity of speed-modulation patterns in sensory neurons

Our next aim was to characterize the gain modulation patterns in more detail. As before, we focus primarily on two-subunit orientation-selective neurons. To analyze joint patterns of speed modulation, we concatenated the modulators of both subunits. We then performed a clustering analysis of such concatenated modulators. Because the speed correlation analysis suggests a continuum of tuning patterns (see Fig.5c-e), our aim was not to identify a specific number of discrete tuning “types” but rather to summarize their diversity and reveal general trends in the population. We therefore chose to analyze 12 clusters; however, analyses with different cluster numbers led us to similar conclusions, supporting the existence of a continuum of modulator properties.

Cluster centroids reveal a diversity of speed modulation patterns in the analyzed population (Fig.6a). In agreement with the previous analysis, the average modulator pairs in each cluster consist of one stronger and one weaker average subunit (Fig.6a, darker and lighter shades of the same color, respectively). Strong differences between centroids suggest that neurons in each cluster combine sensory features with different relative weights at different speeds of movement. In some clusters, the difference between the average modulators is very pronounced (e.g., clusters 1, 8 and 9), while in others modulator averages follow a similar trend (e.g. clusters 6, 11 and 12).

Modulator averages are representative of the general trend within each cluster; however, they may not characterize well the modulation patterns in individual neurons. In particular, due to the asymmetry of modulator strengths, the clustering outcome depends more on the stronger subunits. To explore the fine structure of individual clusters, we visualized all modulator pairs, sorted by the center of mass of the weaker modulator (Fig.6b). Within each group, the strongest modulators are relatively similar, as indicated in the weak spread of their centers of mass (Fig.6b, left part of each panel, colored dots). However, weaker modulators are typically diverse even within the same cluster (Fig.6b, right part of each panel, colored dots). Their centers of mass are often broadly distributed spanning the entire range of running speed values (e.g. clusters 2, 3, 4, 8 and 10).

The joint distributions of the stimulus directions preferred by the subunits were similar across the clusters, with individual subunits separated by 180 degrees with the highest probability (Fig. 6 c: histograms corresponding to four largest clusters). We did not observe systematic preferences for stimulus directions within clusters, suggesting that different patterns of speed modulation act on all neurons, regardless of their sensory tuning.

**Figure 6:**
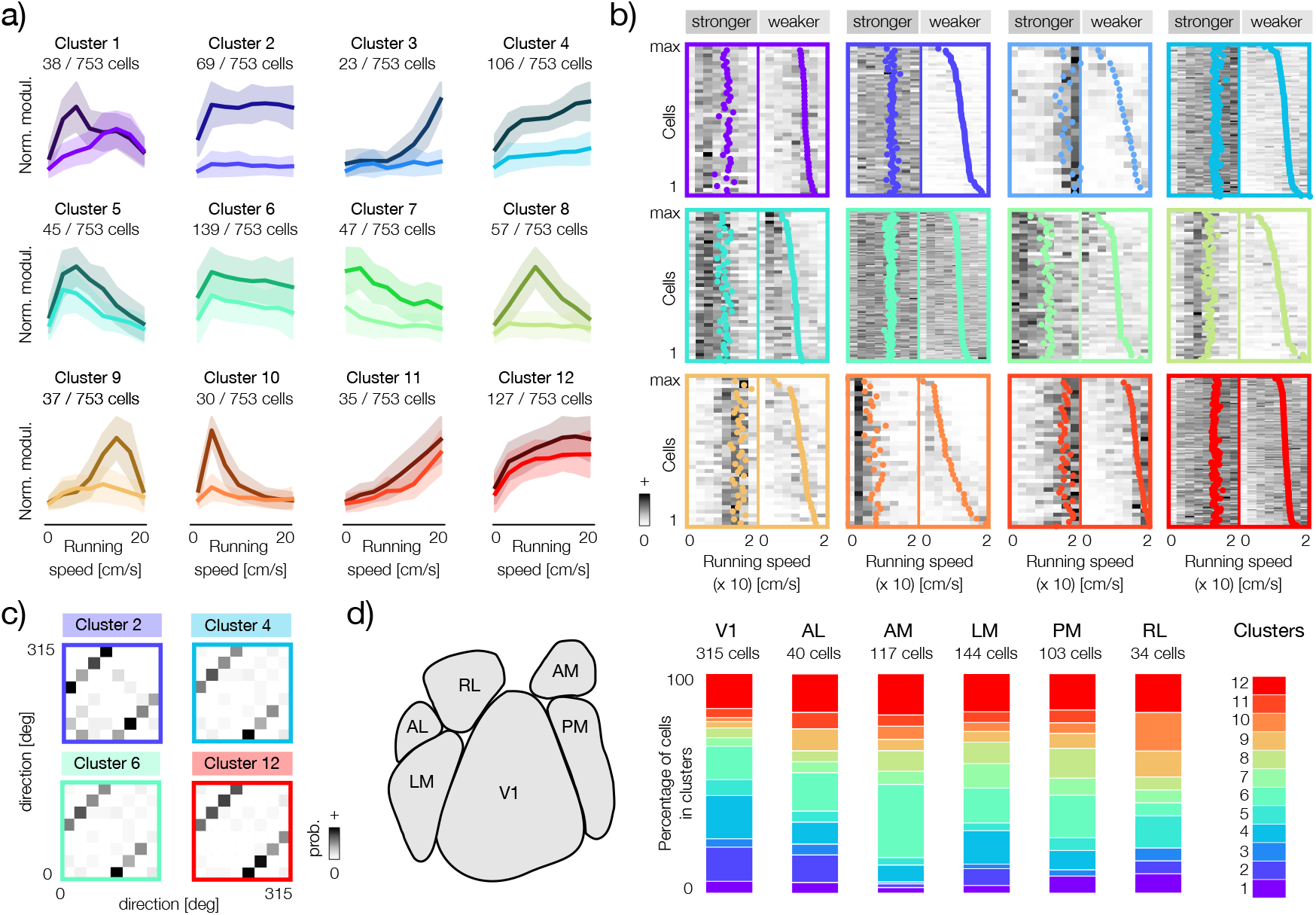
Clustering two-subunit neurons shows diverse effects of locomotion on sensory tuning. **a)** Twelve clusters of speed modulators in neurons with two subunits. Bold lines correspond to cluster means (centroids) and shaded areas to cluster standard deviations. Weaker modulators are plotted with a lighter color variant. **b)** Individual modulators in each of the twelve clusters. Each panel corresponds to an identically positioned panel in a, where colors of centroids in a match the color of frames in b. Each panel consists of two parts corresponding to the stronger and weaker modulator of each cell, separated by a thin, vertical line. Colorful dots correspond to the centers of mass of each modulator. Within each cluster neurons are sorted by the center of mass of the weaker modulator. **c)** Joint distributions of direction selectivity of fitted subunits in four largest clusters. **d)** Left panel: a schematic of the anatomical division of the mouse visual cortex according to the Allen Brain Observatory (replotted from [21]). Right panel: distributions of cluster memberships in the six areas of the visual cortex. Each vertical bar corresponds to one area, the total bar height corresponds to 100 % of neurons in that area (the absolute number of neurons is denoted above each bar). Colors (right panel, color bar) correspond to clusters visualized in panels a and b.

To explore the possibility that certain types of speed modulation are specific to different areas of the brain, we analyzed the probabilities of cluster membership for neurons in the six areas of the visual cortex (Fig.6d). Most of the clusters were represented in all visual areas. Upon closer inspection, certain trends become apparent. In one example, clusters 2 and 4, where modulation saturated rapidly after the onset of locomotion, were more prevalent in the primary visual cortex (V1) than in higher areas. In another example, cluster 6, characterized by a weak decrease in gain at higher speeds, was most strongly represented in the area AM. In general, these results are consistent with the notion that different modulation patterns are broadly distributed throughout the visual cortex. We also note that because of the large differences in the number of neurons analyzed within each area, it is difficult to reach systematic conclusions.

To further characterize the impact of speed on sensory representations, we analyzed the patterns of speed modulation in single-subunit, direction-selective neurons (Fig.7. We grouped neurons into nine clusters that, as previously, we consider to be representative of broader trends, rather than corresponding discrete and isolated types. Clustering revealed various forms of modulation (Fig. 7a). The gain in the largest cluster of neurons (cluster 1) saturated rapidly after the onset of movement. In other clusters, gain increases monotonically with speed (e.g. clusters 5 and 8) or shows tuning for specific running speeds (clusters 2, 3, 4, 6, 7, and 9). The analysis of sensory tuning within the clusters did not indicate any systematic relationship between the speed modulation pattern and the preferences of the direction of stimulus motion (Fig.7b). To determine whether the prevalence of these groups of neurons is a characteristic of specific anatomical regions, we examined their distribution in the six visual cortical areas (Fig. 7c). The nine modulator clusters were present in all areas, and their relative proportions varied. Although the inter-area differences are subtle, certain trends are clearly visible. Interestingly, the primary visual cortex had the largest fraction of cells whose gain saturates even at minimal running speeds (cluster 1), which resembles a similar trend revealed by the clustering distribution of two-subunit cells (Fig.6d).

**Figure 7:**
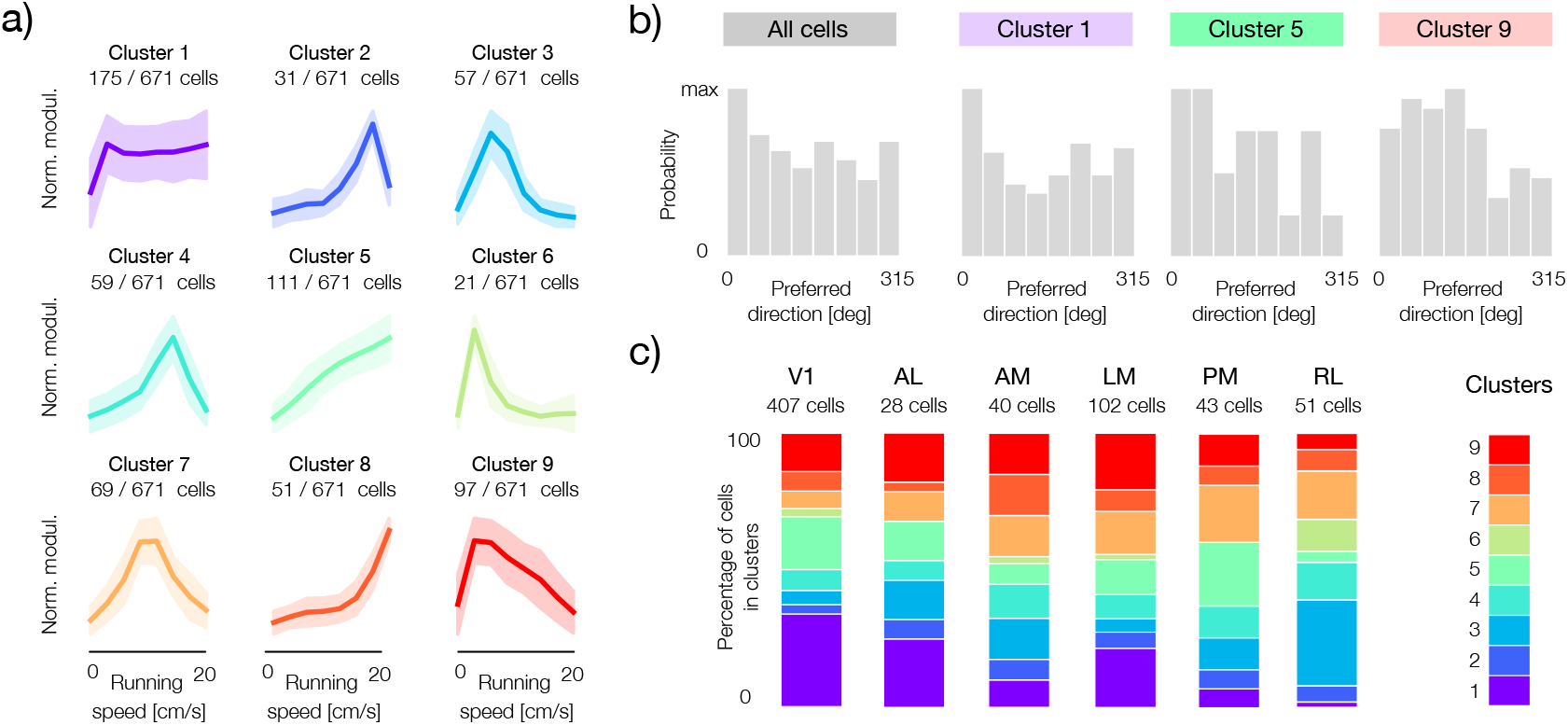
Clustering single-subunit neurons shows variability of running speed tuning. **a)** Nine clusters of speed modulators in neurons fitted with one subunit. Bold lines correspond to cluster means and shaded areas to cluster standard deviations. **b)** Distributions of preferred directions for single-subunit neurons. Distribution across all cells (left panel) and three example clusters (colored), corresponding to panel a). The clusters generally show no clear tuning bias for specific stimulus directions. **c)** Proportional distributions of modulator clusters within six areas of the visual cortex. The colored segments in each vertical bar show the percentage of neurons belonging to each cluster (color bar corresponding to panel a), for each visual area. The total number of neurons in each area is given above the bars.

### Differential behavioral modulation of sensory neurons can support a variety of tasks

Behavioral modulation of sensory tuning likely supports a range of computational tasks. Understanding these computations in detail is, of course, a major challenge that lies outside the scope of the current study. We can, however, consider in more general terms how different types of behavioral modulation of visual neurons can support downstream computations.

Brain areas downstream from the visual cortex can read-out task-specific variables from populations of sensory neurons that combine visual and behavioral signals (Fig. 8a). We note that the performance of any computational task depends on how accurately neurons represent the inputs required for this task. For computations that involve the manipulation of sensory and behavioral signals, populations of neurons should encode these signals with the desired level of accuracy. We illustrate this intuition in Fig. 8b. For example, to achieve high performance in a difficult sensory task (Fig. 8b, left panel, grayscale), the neural population needs to encode sensory signals with an accuracy exceeding a high threshold (Fig. 8b, left panel, vertical blue line). At the same time, the behavioral signal could be represented more coarsely by setting the accuracy threshold lower (Fig. 8b, left panel, horizontal blue line). Similarly, for a difficult behavioral task, behavioral signals should be represented and processed with higher accuracy than the sensory ones (Fig. 8b, middle panel, blue lines). A demanding joint task requires both sensory and behavioral signals to be encoded with high precision (Fig. 8b, right panel). Populations of neurons that simultaneously encode sensory and behavioral inputs with high precision can therefore support a broad range of tasks that require these variables.

**Figure 8:**
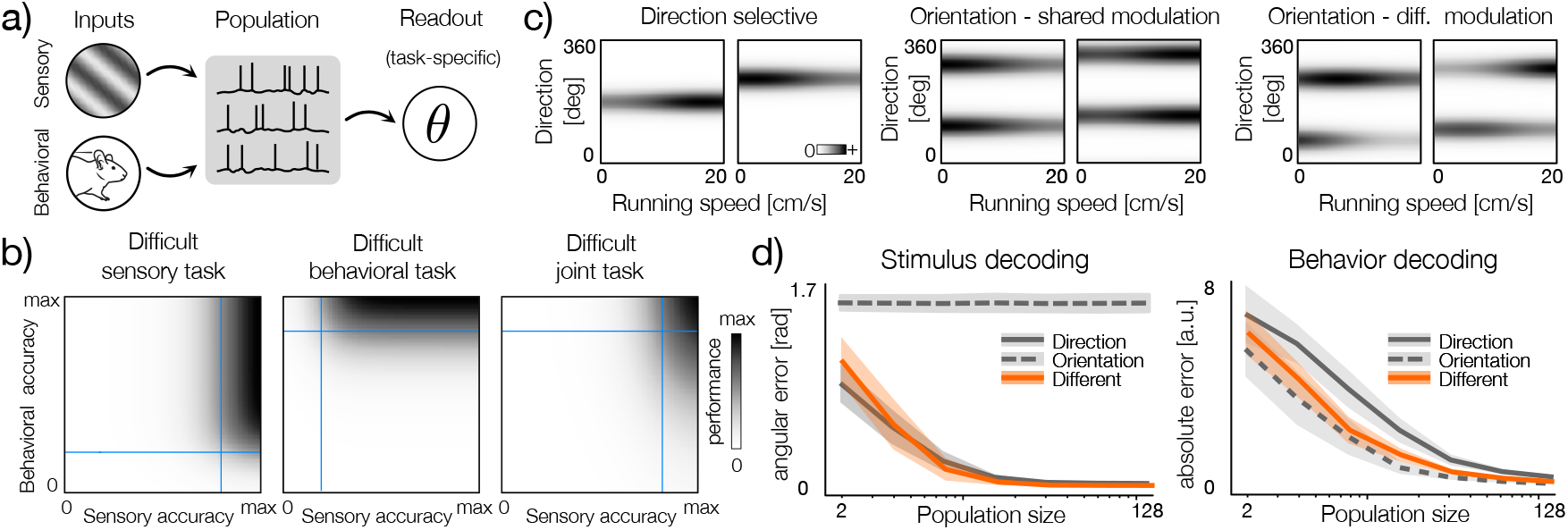
Computational benefits of differential tuning. **a)** Schematic of computation with behavioral and sensory signals. Population of neuron processes the inputs such that down-stream neurons can read-out task-specific quantities (denoted by *θ*). **b)** Performance of hypothetical tasks (grayscale) plotted as a function of the accuracy of representation of behavioral and sensory inputs in the population. Left: a difficult sensory task - to achieve high performance sensory input has to be encoded with high accuracy (vertical blue line) while the behavioral input can be represented with low accuracy (horizontal blue line). Middle: same as left but for a difficult behavioral task. Right: Same as left but for a difficult joint task. **c)** Example tuning curves representative of three different neuron types modulated by running speed. Left: direction selective neurons. Middle: orientation selective neurons with shared modulation of subunits and symmetric tuning curves. Right: orientation selective neurons with differently modulated subunits. **d)** Left: simulated stimulus decoding from populations of different neuron models. Lines denote errors averaged over multiple random populations and shaded bars denote the standard deviation of the error. Direction-selective cells (gray solid line and shaded area) and differently modulated orientation-selective neurons (orange line and shaded area) models have very similar performance. Orientation-selective neurons with shared modulation (gray dashed line and gray are) has the constant average error of approximately *π/*2 regardless of population size. Bottom: simulated decoding of the behavioral variable that modulates neuronal activity. Colors and markers the same as above. Direction-selective neurons yield highest decoding error while multiple both types of orientation-selective neurons models show comparable, better performance.

**Figure 9:**
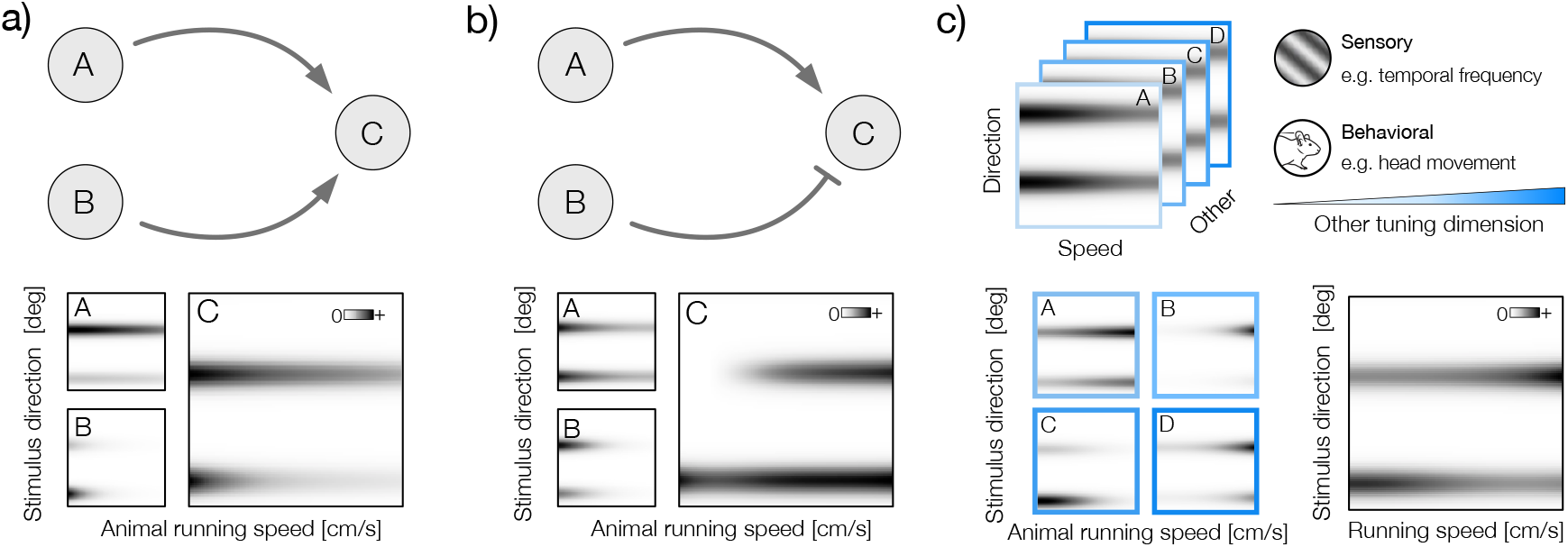
Hypothetical mechanisms of differential modulation. **a)** Top: a schematic of excitatory convergence of two neurons (A and B) on the third neuron (C). Bottom: Example direction-speed tuning curves of neurons A and B (left, smaller panels) - subunits of both neurons share a common speed modulator. Resulting tuning curve of the neuron C (right) shows differential tuning. **b)** Same as a but for an example excitatory-inhibitory convergence of two orientation-selective neurons with shared modulation of subunits. **c)** Schematic of a complex tuning. Top: a 3D tuning curve - in addition to direction and speed, the neuron is tuned to a third variable. This other variable could be either sensory (e.g. temporal frequency), or behavioral (e.g. head movements). Different values of the additional variable are denoted by letters A-D. Bottom: direction-speed tuning changes with the third tuning variable (left; small panels, blue frames). Right: the resulting direction-speed tuning curve shows differential modulation.

To understand how different types of speed-induced modulation in the visual cortex can support the precision of sensory and behavioral representations, we simulated neural populations and performed a decoding analysis. We considered three types of neurons: direction selective, orientation selective with shared behavioral modulation, and orientation selective with different modulation of subunits (Fig.8c). We assumed that a downstream area randomly pools neurons of a given type and simulated decoding from multiple random populations (see Methods). This simple approach has a number of advantages. First, it allows us to study the encoding performance of a given neuron model without explicitly optimizing the population. We thus focus on distributions of solutions instead of just a single optimum. Second, it allows us to study how well behavioral and sensory signals are encoded by different neural architectures without reference to the specific computational task that those neurons may perform. The only assumption is that these tasks require an accurate representation of the stimuli and the behavioral state. Third, random grouping requires less fine-tuning, and the nervous system could easily implement such a pooling scheme [24, 25].

As expected, we found that direction-selective, single-subunit neurons encode sensory stimulus direction with the highest accuracy; the average error decreases most rapidly as a function of the population size (Fig. 8d, left panel, solid gray line). In contrast, behavior-modulated orientation selective neurons yield the highest average error in decoding stimulus direction (Fig. 1d, left panel, gray dashed line). This high error level is, of course, due to the ambiguity in orientation selective cells that do not distinguish between stimulus directions rotated by 180 degrees. Relative performance reverses when decoding the modulating behavioral variable (Fig. 8d, right panel). Here, orientation selective neurons represent the behavioral variable with the highest accuracy at a given population size. Direction-selective neurons typically require twice as many neurons to achieve the same level of decoding error. This is because, in order for behavioral signals to be encoded at all, they must coincide with a stimulus represented by a neuron. Because orientation-selective neurons, by definition, have broader tuning than direction-selective ones, they respond more frequently and thus encode behavior with higher accuracy. In the supplemental material, we present results of a similar analysis, where neural activity was simulated with real tuning curves (Supp. Fig. 2).

Simulated decoding analysis demonstrates that populations of direction- and orientation-specific neurons trade off the accuracy of encoding stimulus direction and modulating behavioral signals. Interestingly, the neuron model in which subunits are modulated differently by behavior navigates this trade-off and achieves good performance for both variables (Fig. 8d, orange line). During the encoding of stimulus direction, behavioral modulation differentiates responses to stimulus directions encoded by individual subunits, thus resolving the directional ambiguity present in orientation-tuned cells. The resulting decoding performance is very close to that of direction-selective neurons. In a sense, differential behavioral modulation allows for the read-out of the stimulus direction from orientation-selective neurons. If a sensory task requires resolving the direction of visual motion and not just it’s orientation, this mechanism can effectively increase the “resolution” of the sensory code. At the same time, because independently modulated orientation-selective neurons combine more than one stimulus direction, they encode behavioral inputs with higher accuracy than populations of direction-selective cells of the same size. It is therefore possible that behavioral modulation of sensory tuning can support a broad range of computational tasks that require accurate representation of sensory and behavioral information. We note, however, that readout mechanisms different from the one considered here may impact the relative benefit of differential modulation.

### Possible mechanisms of differential modulation of sensory neurons in the visual cortex

Modulation of sensory tuning by behavior can be instantiated by many possible underlying mechanisms. Here, we discuss examples of how such tuning could be implemented in simple neuronal circuits. First, we observe that the emergence of complex selectivity in sensory systems can often be explained by combinations of simpler sensory features. A classical example is the formation of phase-invariance in complex cells by pooling the outputs of simple cells [26]. Behavioral changes in sensory tuning could also be achieved by combining the outputs of cells with a simpler form of tuning (Fig.9a-b). For example, the excitatory convergence of two orientation-selective neurons, whose gains are differentially modulated by speed, can easily generate a complex form of tuning in the downstream cell (Fig.9a). Due to the dependency of input strength on locomotion speed, the downstream cell will modulate its gain to one of the stimulus directions differently than to the other. A specific case of this scenario is the convergence of two direction-selective cells whose gains are modulated differently by running speed. Another way to implement differential modulation is through excitatory-inhibitory convergence (Fig.9b). Here, one of the upstream cells suppresses the downstream one, generating a complex form of sensory-behavioral tuning. These effects can be potentiated if upstream cells exhibit “asymmetric” orientation tuning, where the response strength to one of the stimulus directions is stronger, or if downstream cells combine the outputs of more than one upstream neuron. Because differential modulation can arise in such simple convergent motifs, it is also possible that it is merely a consequence of circuitry in the visual cortex rather than a manifestation of a purposeful computational strategy. The combination of simpler features to implement more complex tuning could also, in principle, be accomplished by a single neuron that exploits dendritic nonlinearities.

In addition to stimulus direction and speed, neurons in the visual cortex are tuned to many other sensory and behavioral variables, such as temporal frequency or head movements [27]. A complex form of tuning is possible, where an additional variable impacts the interaction between the encoding of running speed and stimulus direction (Fig.9c). The resulting direction-speed tuning curve, averaged across this additional variable, could show changes in direction preference at different locomotion speeds. Such a mechanism could be an alternative form of a complex interaction between sensory and behavioral variables that goes beyond mere gain modulation during movement.

## DISCUSSION

In this work, we provide evidence that movement can modify the sensory tuning of visual neurons in subtle ways. This modification takes a specific form: instead of modulating the gain of the neuron’s responses to all stimuli, running speed modulates the neuronal gain depending on the stimulus value. Responses to stimuli represented by different functional subunits may therefore be enhanced or suppressed in various ways during behavior.

The functional role, or most likely multiple roles, of this modulation remains a mystery. We point out, however, it’s two computational benefits. First, different responses to stimulus directions enable the readout of this feature even from populations of orientation selective cells. Orientation selectivity is a form of invariance or compression; it ignores the direction of stimulus movement. A mechanism that breaks the symmetry between movement directions separated by 180 degrees effectively removes this ambiguity. Here, we discuss behavioral gain modulation as one possible mechanism. We note, however, that even when neglecting the impact of behavioral state, orientation selective neurons can respond with different strengths to stimuli at two preferred directions, which is manifested in “bimodal” direction-tuning curves, where each mode has different amplitude [23]. This mechanism is wide-spread in the mouse visual cortex [22]. We consider it possible that this property is inherited from the differential behavioral modulation of direction-selective subunits.

The second computational benefit is the accurate multiplexing of the modulating behavioral variable in a sensory population. This mechanism enables the visual cortex to maintain information about the sensory environment and the behavior of the organism. These two types of signals are very likely required to support computations that drive behavior. While we do not know the specific nature of these computations, several examples have already been discussed in the field. A recent study has suggested an elegant example by demonstrating that joint tuning to running speed and spatial frequency may support depth estimation via motion parallax [28]. Other well-known examples include the signaling of sensorimotor mismatches [29] or the estimation of the direction of movement from visual and proprioceptive signals [9]. We believe that many types of not-yet-described computations in the visual cortex may require the simultaneous, joint processing of behavioral and sensory signals using the mechanisms described here. Importantly, these benefits are accessible even with randomly pooled populations. This finding is in agreement with studies that demonstrate the utility of heterogeneity, variability, and sometimes even randomness of tuning in sensory representations and neural circuits [24, 25, 30, 31].

An open set of questions concerns the biological implementation of gain modulation by running speed at the single-subunit level. While we discussed possible implementations of this form of modulation, we do not consider our discussion exhaustive. Multiple studies have demonstrated that the selectivity of neurons in the visual cortex cannot be easily captured by a single linear receptive field and that even individual neurons combine multiple stimulus features, often referred to as subunits [13, 16, 19, 32]. Given how widespread subunit architectures are in the visual system, we consider it likely that behavior can influence individual subunits differently, giving rise to the effects discussed here. Whether they are inherited from the tuning of upstream neurons, are a manifestation of complex circuit interactions, or even the computational capabilities of individual cells remains an open question.

Another important question concerns multiple gain modulators in the visual cortex. Gain modulation is a prominent mechanism that contributes to neuronal variability in behaving animals [33, 34]. Single neurons can be subject to multiple internal states that affect the gain of their sensory responses [33, 35, 36]. It is possible that some of the speed-induced modulation reported here is caused not directly by running speed, but by other correlated factors, e.g., attentional state, arousal, or head movements. Although we cannot separate those factors with complete certainty, our results indicate that the sensory tuning of individual neurons is dynamically modulated in behaving animals, regardless of the origin of the modulatory signal.

The current results complement our recent study, in which we demonstrated that gain modulation by locomotion is required to maintain an accurate and efficient sensory representation [37]. The theory presented in [37] predicts that gain modulation optimized to maintain coding accuracy takes a specific form: the strength of sensory responses should increase rapidly at the start of locomotion and then saturate. Interestingly, qualitatively similar speed modulation patterns are identified by clustering analyses in this study (Fig.7 a, cluster 1 and Fig.6a, cluster 2 and 4). Neurons whose subunits are modulated in a way most consistent with this theory are mostly abundant in the primary visual cortex. These results enable us to speculate that gain saturation is a signature of a “sensor-like” neuron whose role is to encode and transmit sensory information accurately across behavioral states. Other patterns of gain modulation may indicate different types of computation. Exploring these ideas further remains, of course, a subject of future work.

Addressing all of these questions in detail will require dedicated experimental work. A particular challenge in characterizing neuronal tuning is gathering comparable amounts of data in different behavioral states, which change independently of the experimenter. In one example, we envision dedicated experiments where temporal and spatial frequencies are varied systematically in addition to stimulus direction. Understanding neuronal tuning to these additional sensory features could provide much deeper insight into the mechanisms of behavioral modulation in sensory systems. To gather sufficient amounts of data during time-limited experiments, the application of closed-loop experimental design methods presents particular promise (see e.g. [38]). We also foresee the possibility of answering these questions through the application of dedicated statistical models that can separate different sources of variation in neuronal activity (e.g. [33, 39, 40]).

## ACKNOWLEDGEMENTS

We thank Laura Busse, Lukas Meyerolbersleben and Ann Hermundstad for helpful discussions.

## METHODS

### Simulated populations of gain-modulated neurons

We simulated neurons tuned to an angular variable *ϕ* ∈ [ − *π, π*] (e.g. stimulus direction) using von-Mises-like tuning curves of the form: *r*(*ϕ*) = exp(*κ* ∗ cos(*ϕ*_*i*_ − *ϕ*)), where *ϕ*_*i*_ is the preferred tuning direction and *κ* = 4 controls width of the tuning function. Each sensory tuning was normalized so that the maximal response was equal to 1. We simulated the speed tuning by combining 8 radial basis functions of the following form 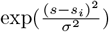 where *s*_*i*_ denotes the position of the peak and *σ*^2^ = 10 is the width of the basis function. The peak positions were linearly spaced on the [0, 20] interval. Each speed modulator *m*(*s*) was expressed as a linear combination of basis functions: 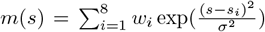 where the weights *w*_i_ were randomly drawn from a Gaussian distribution with standard deviation equal to 0.5 To generate the response *r* to a particular stimulus direction *ϕ* and speed *s* the speed modulator was applied multiplicatively after exponentiation. The functional form of the response took the form: *r*(*s, ϕ*) = *r*(*ϕ*) exp(*m*(*s*)) + *η*, where *η* is the additive Gaussian noise with standard deviation *σ* = 0.2. For two-subunit model neurons with shared modulation, we generated the same modulator for both subunits, and the neural response was summed over both subunits. For two-subunit neurons with independent modulation, separate modulators were randomly generated for each subunit. To simulate asymmetry of the modulator strength, a random number uniformly distributed on the [0, 0.2] interval was subtracted from the weights *w*_*i*_ of the weaker subunit.

We simulated 500 random populations for the following population sizes: 2, 4, 8, 16, 32, 64, 128 neurons. In each of the simulations we generated 500 random stimuli *ϕ* uniformly distributed on a circle and speed values *s* uniformly distributed on the [0, 20] interval.

We performed the maximum-a-posteriori (MAP) decoding. First, we computed the joint posterior over the stimulus *ϕ*_*t*_ and speed *s*_*t*_ given the noisy response of the population 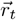, i.e. 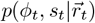. We then marginalized the joint posterior to generate two marginal posteriors 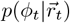 and 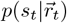. The maxima of these posteriors were taken to be the decoded estimates 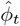 and *ŝ*_*t*_, respectively. All computations were performed numerically, using discretized values of parameters. Because the prior distributions *p*(*ϕ*_*t*_), *p*(*s*_*t*_) were both uniform, this procedure was equivalent to maximum likelihood (ML) decoding. To quantify the speed decoding error we used the squared distance averaged over samples, i.e. ⟨(*s*_*t*_ − *ŝ*_*t*_)^2^⟩_*t*_. To quantify the stimulus direction decoding error, we used the average angular distance 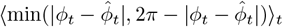.

### Simulated decoding analyses

For the analysis displayed in the Supplemental Material (Fig.S2), we performed the MAP decoding identically as with the simulated tuning curves (see above). We used speed-orientation tuning curves from three areas with the highest number of retained neurons. We used only the two-subunit cells where subunits were separated by 180 degrees (numbers of cells per area are stated in Supp. Fig. 2d-f). For each area, we generated populations of the following sizes: 2, 4, 8, 16, 32, 64, 128. For each size value, we generated 200 random populations, each sampled from tuning curves of a corresponding area with replacement. During simulation, each of the populations was used to decode the random stimulus and speed values 1000. Stimulus values corresponded to the eight directions used in experiments, while speed values corresponded to the speed-binning used to derive tuning curves. Each tuning curve was normalized so that its maximum was equal to 1. Neural responses were generated by adding a random Gaussian number with zero mean and standard deviation *σ* = 0.2 to the value of the tuning curve corresponding to the speed and orientation pair. To generate neurons that are data-matched but share modulation across the subunits, we replaced modulators of each of the subunits with their average.

### Data preprocessing

We used data made available by the Allen Brain Observatory [41]. The datasets comprised of two-photon calcium imaging of neural populations recorded in behaving mice. The animals were exposed to drifting grating stimuli at cpd spatial frequency, at 8 different orientations linearly spaced 45 degrees apart between 0 and 360 degrees and 5 temporal frequencies (1, 2, 4, 8, 15 Hz).

To reduce the variability of the data and focus the analysis on variation above the baseline, we thresholded the ΔF/F signal by setting all negative values to zero. Following thresholding, we normalized the activity of each cell by dividing it by its standard deviation.

To ensure the robustness of our conclusions, we applied several data filtering steps. We included datasets where no direction-speed bins were empty (48 % of all cells), joint tuning curves were significantly well-fitted by our sparse decomposition model (26 % of cells with non-empty bins), cells exhibited sensory tuning (81 % of cells with significant fit), cells whose modulation along the running speed axis was significant (87 % of cells with sensory tuning). Finally, we aimed to minimize the distortions in estimated tuning due to differences in the amount of data per bin and disparities between the range of temporal frequencies presented for each bin. To achieve this, we excluded any datasets with less than 1 second of data in any of the 64 bins of direction-speed tuning curves. Additionally, we removed datasets where more than 16 of the 64 bins were derived from averages that included fewer than 3 different temporal frequencies (out of 5 total). Following all selection criteria, we obtained a dataset of 1693 neurons from 100 recording sessions.

### Computation of tuning curves

We divided the running speed into eight linearly spaced bins that ranged from 0 and 20 cm/s. Individual mice revealed different running preferences and generated different distributions of the speed across recorded sessions. The considered speed binning allowed for consistent comparisons across different datasets with varied velocity distributions. Subsequently, we binned the transformed ΔF/F values of each cell according to the eight orientations, five temporal frequencies and eight running speed bins. The activity within each bin was then averaged to obtain the average response *r*_*o,v,f*_ where *o, v, f* denote the stimulus orientation, running speed and the stimulus temporal frequency, respectively. We refer to these average activity tensors as 3D tuning curves. Averaging along the frequency dimension we obtained *r*_*o,v*_, which we refer to as direction-speed tuning curve.

### Tuning specificity

To assess the sensory tuning specificity of each cell, we computed their stimulus tuning curves *R*^*t,s*^ by averaging each direction-speed tuning curve *R*^*t*^ along the speed dimension. We introduce the normalized entropy of each tuning curve 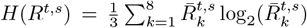, where 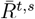 is the sensory tuning curve normalized, such that its entries sum up to unity. The normalized entropy *H* is equal to 0 if the tuning curve is uniform and to 1 if the neuron responds to only one stimulus. We quantified the tuning specificity of each neuron as *S* = 1 − *H*(*R*^*t,s*^). Neurons were classified as stimulus selective and included in further analysis, if their sharpness values exceeded the 95th percentile of the null distribution derived from the set of randomly shuffled tuning curves (see the description of goodness of fit assessment).

### Bootstrap test of speed modulation

We performed a bootstrap test to test for the significance of speed modulation. For each neuron, we generated 1,000 3D tuning curve tensors by resampling the data that contributed to each bin (defined by stimulus orientation, stimulus temporal frequency, and running speed). For each of the bins and in each of the 1000 random tuning curves, we resampled identical number of data points, corresponding to the responses of the neuron to the same stimulus direction and temporal frequency, but to random speed values. This procedure ensured that the resulting tuning curves were velocity-invariant while preserving the direction and frequency tuning properties of the neuron, as well as the number of data samples in each bin. Each of the randomly resampled 3D tuning curves was averaged across the temporal frequency dimension, as with the original data.

We used the following test statistic to assess the modulation strength: 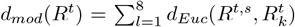, where *R*^*t,s*^ is the sensory tuning curve (see above), 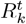 are sensory tuning curves at each speed bin *k* and *d*_*Euc*_ is the Euclidean distance. Modulation of sensory tuning by running speed was considered significant if the value of the test statistic evaluated for the real tuning curve exceeded the 95th percentile of the null distribution of the test statistic generated by bootstrapping for the considered cell.

### Bootstrap test of modulator difference

First, for each neuron with two significant subunits we performed a bootstrap procedure similar to the one described above. The procedure entailed resampling the data that contributed to each tuning curve bin (defined by stimulus orientation, stimulus temporal frequency, and running speed). For each cell we averaged two fitted modulators to compute a simulated shared modulator. Samples within each bin of the bootstrapped tuning curves were multiplied by this simulated shared modulator. This procedure ensured that resulting tuning curves preserved the direction and frequency tuning properties of the neuron, as well as the number of data samples in each bin. The only differing factor was the shared, multiplicative gain of the significant subunits. We generate 1000 such bootstrap samples for each neuron with two significant subunits.

To compute test-statistics 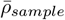, we optimized a non-negative curve (correlation-maximizing fit) that maximized the average correlation with each of the two modulators. For optimization, we used the BFGS algorithm implemented in the minimize function of the scipy package. The distribution of 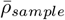 obtained from all bootstrapped tuning curves for individual neurons served as the null distribution for significance tests. For each real neuron, we optimized the correlation maximizing fit in the same way and computed the value of the test statistic 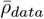.

### The sparse decomposition model

We designed a statistical model to parameterize the patterns of joint tuning to stimulus direction and running speed by decomposing the tuning curves of individual neurons. The model assumes that each neuron pools the responses of direction-selective subunits, where each subunit is multiplicatively modulated by running speed.

A direction-speed tuning curve of the *t*−th neuron *R*^*t*^ is a two-dimensional, *K* matrix, where *L* = 8 is the number of speed bins and *K* = 8 is the number of stimulus direction bins. Each element 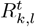 represents the mean response of the neuron *t* to a grating of a specific direction *k* and the speed interval *l*.

Basis functions *ϕ*^*i*^ represent sensory tuning to one of the *K* preferred directions and are effectively zero-matrices with entries in one row set to 1. The total number of basis functions is therefore equal to the number of possible stimulus directions i.e. 8. The model parameters 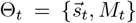 consist of a matrix of modulators *M* (ℝ^*L×*8^), which describe speed-bin-specific modulation, and weights 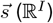, which represent the weight of each orientation-specific subunit. The model reconstructs speed-direction tuning curves as the sum of weighted and modulated basis functions:

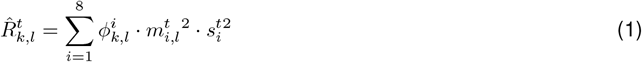

where square is taken to ensure positivity of modulators and basis function coefficients.

We estimated the model parameters for each neuron Θ_*t*_ by minimizing the cost function that combined the mean squared error (MSE) of the reconstruction between the reconstructed and actual data and regularization terms that penalized the absolute value of the subunit coefficients and modulators:

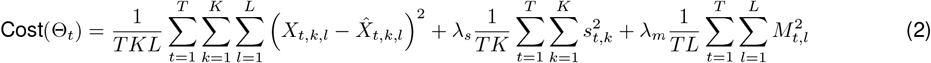

where *λ*_*s*_ and *λ*_*m*_ are the sparsity parameters for subunit coefficients and modulators, respectively. For simplicity we set them to be identical i.e. *λ* = *λ*_*s*_ = *λ*_*m*_. We optimized the model for each neuron using 10 sparsity values, evenly spaced within the range [0.01, 0.1].

For each neuron, the model was fitted using 300 gradient steps, starting from five independent random initializations of parameters. We selected the fits that generate the highest signal-to-noise ratio (SNR) value.

Following the model fitting we multiplied each modulator *m* by the corresponding subunit coefficient *s* (we refer to those as scaled modulators). We found that separating these quantities during fitting resulted in better matches to the data structure; however, following the fit the separation was not useful. For each neuron, we kept only the subunits whose average value of the corresponding scaled modulators was equal to or higher than 5% of the maximal one. We refer to those subunits as “significant subunits”.

### Goodness of fit assessment

For each analyzed dataset, we generated a null set of tuning curves. We did so by randomly sampling the direction-speed tuning curves of 10,000 neurons with replacement. The positions of the 64 bins of each joint tuning curve were then randomly permuted. We fitted the model to each random tuning curve using the procedure described above. For each considered value of the sparsity constraint *λ*, we retained the distribution of reconstruction SNRs obtained in that way. Neurons passed the goodness-of-fit test if their reconstruction quality exceeded the 95th percentile of the null distribution. For each neuron, we selected a single reconstruction corresponding to the lowest *λ* value at which its fit first exceeded the significance threshold.

### Analyses of two-subunit neurons

For each neuron with two significant subunits, we computed the log_2_ of the ratio of the average stronger modulator to the average weaker modulator (Fig. 5b).

To construct null distributions of modulator correlations and modulator-speed correlations, we randomly permuted individual modulators in each neuron across the speed dimension.

To simulate joint modulation of both subunits, we computed the average of both modulators for each neuron. In the next step, we replaced individual modulators in each neuron with this average normalized such that the norm of each subunit was preserved. We added Gaussian noise with standard deviation *σ* = 0.01.

### Clustering analyses

To identify patterns of velocity modulation, we performed K-means clustering on the shapes of the subunit modulators from all neurons that passed the filtering criteria. For each neuron with two significant subunits, we concatenated the scaled modulators (with the stronger subunit leading), resulting in a 16 single feature vector. The feature vectors were divided by their *L*_2_ norm. The number of clusters was chosen heuristically and was equal to 9 of one-subunit neurons and 12 for two-subunit neurons.

## SUPPLEMENTAL FIGURES

**Figure S1:**
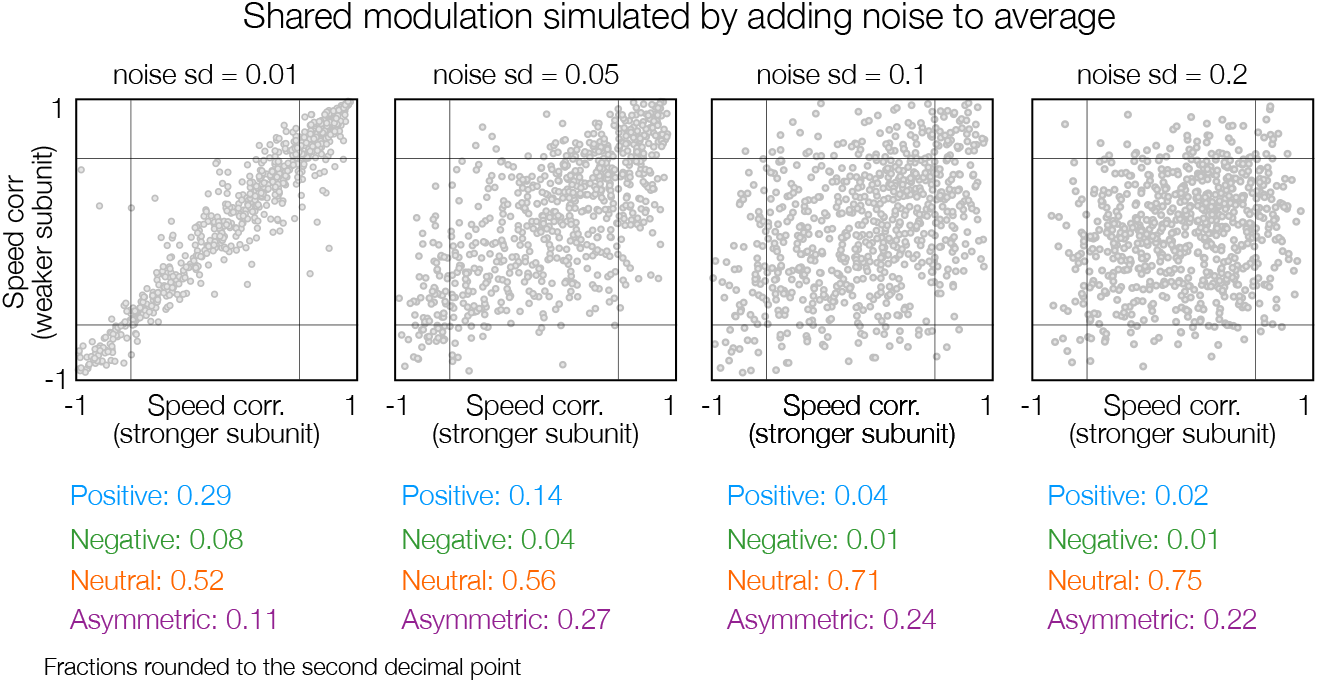
Alternative simulation of shared speed modulation by averaging subunit modulators and adding noise. Top: Scatter plots of correlations of weaker and stronger modulators with the running speed. Each gray circle corresponds to an individual neuron with two significant subunits. Simulations were obtained by averaging modulators of both subunits and adding independent Gaussian noise with three different standard deviations (from the left: 0.01, 0.05, 0.1 and 0.2). Bottom: fractions of neurons in different areas of the plane. Colors correspond to convention in Fig. 5. Numbers are rounded to the second decimal point.

**Figure S2:**
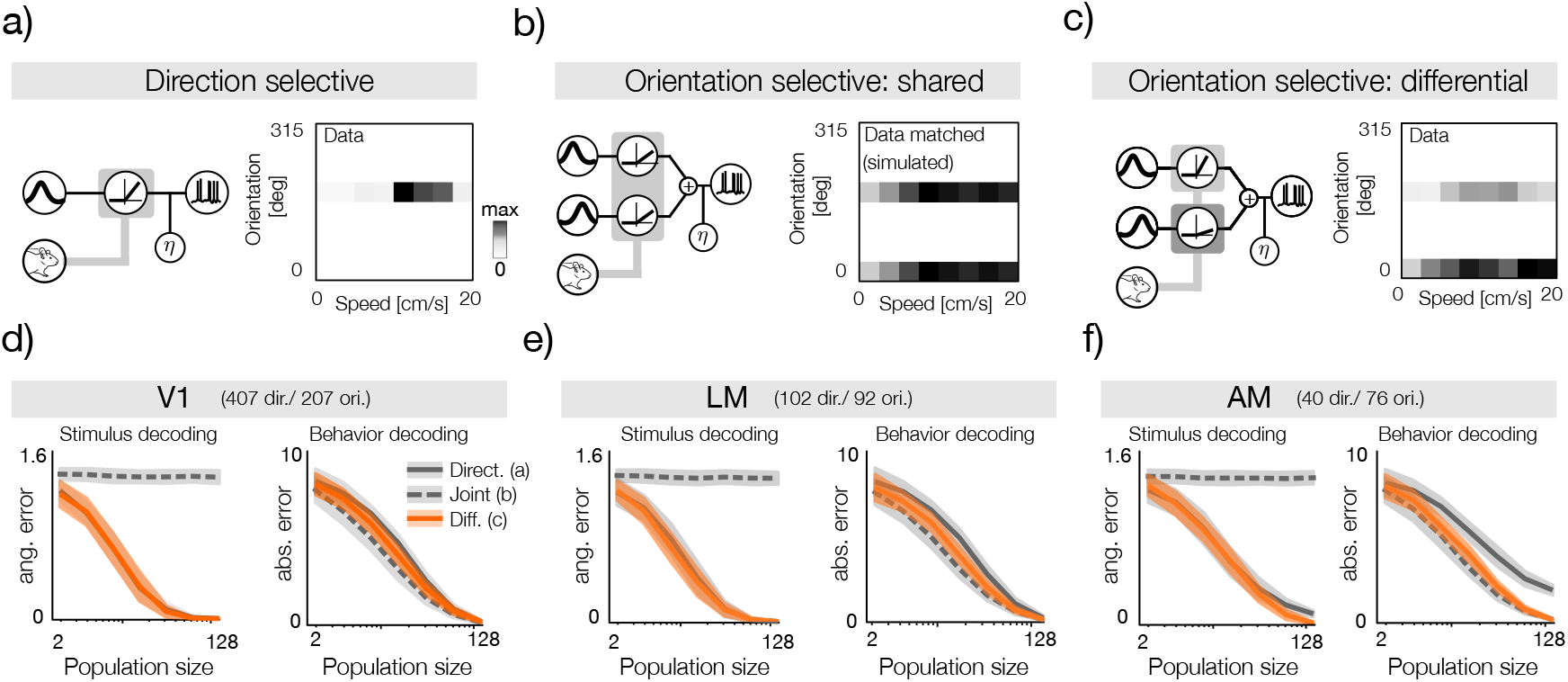
Simulated decoding is consistent with multiplexing of behavioral signals in sensory representations. **a)** A schematic of a direction-selective speed-modulated neuron (left) with an example, corresponding tuning curve fit. **b)** A schematic of an orientation-selective neuron with shared speed modulation of subunits (left) and an example corresponding tuning curve. To ensure identical modulation of both subunits, the tuning curve has been matched to c to ensure identical speed tuning. **c)** A schematic of an orientation-selective neuron with shared speed modulation of subunits (left) and an example corresponding tuning curve fit. **d)** Decoding of the stimulus direction (left panel) and running speed (behavior, right panel) using activity simulated from tuning curves of the primary visual cortex (V1; see Methods for details). In each panel the decoding error is plotted as a function of the population size. Lines correspond to average errors, shaded areas to standard deviations. Populations of direction selective neurons, orientation selective neurons with shared modulation and orientation selective neurons with differential modulation are denoted by gray solid, gray dashed and orange lines respectively. Note that for stimulus decoding (left panel) the orange and solid gray lines and areas overlap almost entirely. **e)** Same as d, but for the area LM. **f)** Same as d, but for the area AM.

